# Identification of a promiscuous conserved CTL epitope within the SARS-CoV-2 spike protein

**DOI:** 10.1101/2021.11.21.469172

**Authors:** Sheng Jiang, Shuting Wu, Gan Zhao, Yue He, Xinrong Guo, Zhiyu Zhang, Jiawang Hou, Yuan Ding, Alex Cheng, Bin Wang

**Affiliations:** Key Laboratory of Medical Molecular Virology (MOE/NHC/CAMS), School of Basic Medical Sciences, Shanghai Medical College(SHMC), Fudan University; National Clinical Research Center for Aging and Medicine, Huashan Hospital, Fudan University, Shanghai, China; Advaccine Biopharmaceutics (Suzhou) Co. LTD, Jiangsu Province, China; Colby College, Waterville, Maine, USA

## Abstract

The COVID-19 disease caused by infection with SARS-CoV-2 and its variants is devastating to the global public health and economy. To date, over a hundred COVID-19 vaccines are known to be under development and the few that have been approved to fight the disease are using the spike protein as the primary target antigen. Although virus neutralizing epitopes are mainly located within the RBD of the spike protein, the presence of T cell epitopes, particularly the CTL epitopes that are likely to be needed for killing infected cells, has received comparatively little attention. In this study, we predicted several potential T cell epitopes with web-based analytic tools, and narrowed them down from several potential MHC-I and MHC-II epitopes by ELIspot and cytolytic assays to a conserved MHC-I epitope. The epitope is highly conserved in current viral variants including the most recent Omicron and compatible with presentation by most HLA alleles worldwide. In conclusion, we identified a CTL epitope suitable for evaluating the CD8+ T cell-mediated cellular response and potentially for addition into future COVID-19 vaccine candidates to maximize CTL responses against SARS-CoV-2.

## INTRODUCTION

Severe Acute Respiratory Syndrome Coronavirus 2 (SARS-CoV-2) was first identified in Wuhan in early 2020, spread at unprecedented speed, and became a disaster to human beings worldwide^1^. Effective vaccines, antiviral drugs, and treatments have high priorities to defend against such challenges. SARS-CoV-2 has four main structural proteins: the envelope, membrane, nucleocapsid, and spike protein that can be considered for inclusion in vaccines. The spike protein has a receptor-binding domain (RBD) that specifically binds to human angiotensin-converting enzyme 2 (hACE2) as a receptor and mediates virus entry into the host cell^2,3^. Neutralizing antibodies recognizing the RBD can block the spike protein from binding to the hACE2 and inhibit virus entry^4,5^. Therefore, spike protein has been the primary choice as the immunogen in candidate vaccines.

Although protection against disease via vaccine-induced neutralizing antibodies has been demonstrated, the elimination of SARS-CoV-2 infection within the host is also essential. The numbers of mild and asymptomatic cases are rising dramatically in recent years and such cases remain infective, prolonging viral dissemination. To eliminate the viral infection, induction of a potent antigen-specific CD8+ T cell response by vaccination is probably critical. To activate a viral-specific CD8+ T cell response, the vaccine must contain highly active major histocompatibility complex class I (MHC-I) epitopes that can be presented by MHC-I molecules to interact with CD8+ T cell receptors (TCR). The potentiation of viral-specific CD8+ T cell responses is dependent upon the high affinity and avidity of MHC-I and TCR binding. There is a lack of information on the CD8+ T cell-recognized epitopes within the spike antigen; consequently, only overlapping peptide pools that covered the whole region of spike antigen have been used routinely to evaluate cell-mediated immunity (CMI) of a vaccine candidate^6,7,8^. However, a few reports had suggested that CTL epitopes are present within the spike protein^9^, but no detailed mapping information had been reported or potential sequences discovered. The identification of those CD8+ T cell epitopes would provide an important tool to evaluate the T cell immunity in vaccinated individuals or patients and was undertaken here.

This study utilized web-based tools to analyze the potentials for transportation associated with antigen processing (TAP) in the human MHC-I epitopes that were predicted by the Immune Epitope Database analysis (IEDB) resource^10^ to be present in peptide pools covering the n-terminal domain (NTD) and receptor-binding domain (RBD) of the spike protein. We demonstrated that peptide 2 (YYVGYLQPRTFLLKY), although it did not give the highest score in the web-based analysis of immunogenicity, was the best epitope for inducing a robust antigen-specific IFN-γ producing CD8+ T response as defined by ELIspot assay. This epitope sequence is also highly conserved among currently discovered SARS-CoV-2 variants.

## MATERIALS AND METHODS

### Mice

Female BALB/c mice (6-8 weeks of age) were purchased from Beijing Vital Laboratory Animal Technology Co., Ltd. (Beijing, China) and Shanghai Jiesjie Laboratory Animal Co., Ltd. (Shanghai, China), and were kept in SPF conditions. All animal experiments were approved by the Experimental Animals Committee of SHMC, and all methods were carried out in accordance with relevant guidelines and regulations. This study was carried out in compliance with the ARRIVE guidelines. All mice was sacrificed after experiment under euthanasia with Isoflurane overdose.

### Peptide pool derived from SARS-CoV-2 Spike protein

The spike receptor-binding domain (RBD) peptide pool (SARS-CoV-2 spike protein 258-518aa) published previously^7^ was used for the study (Table 1).

**Table 1.**
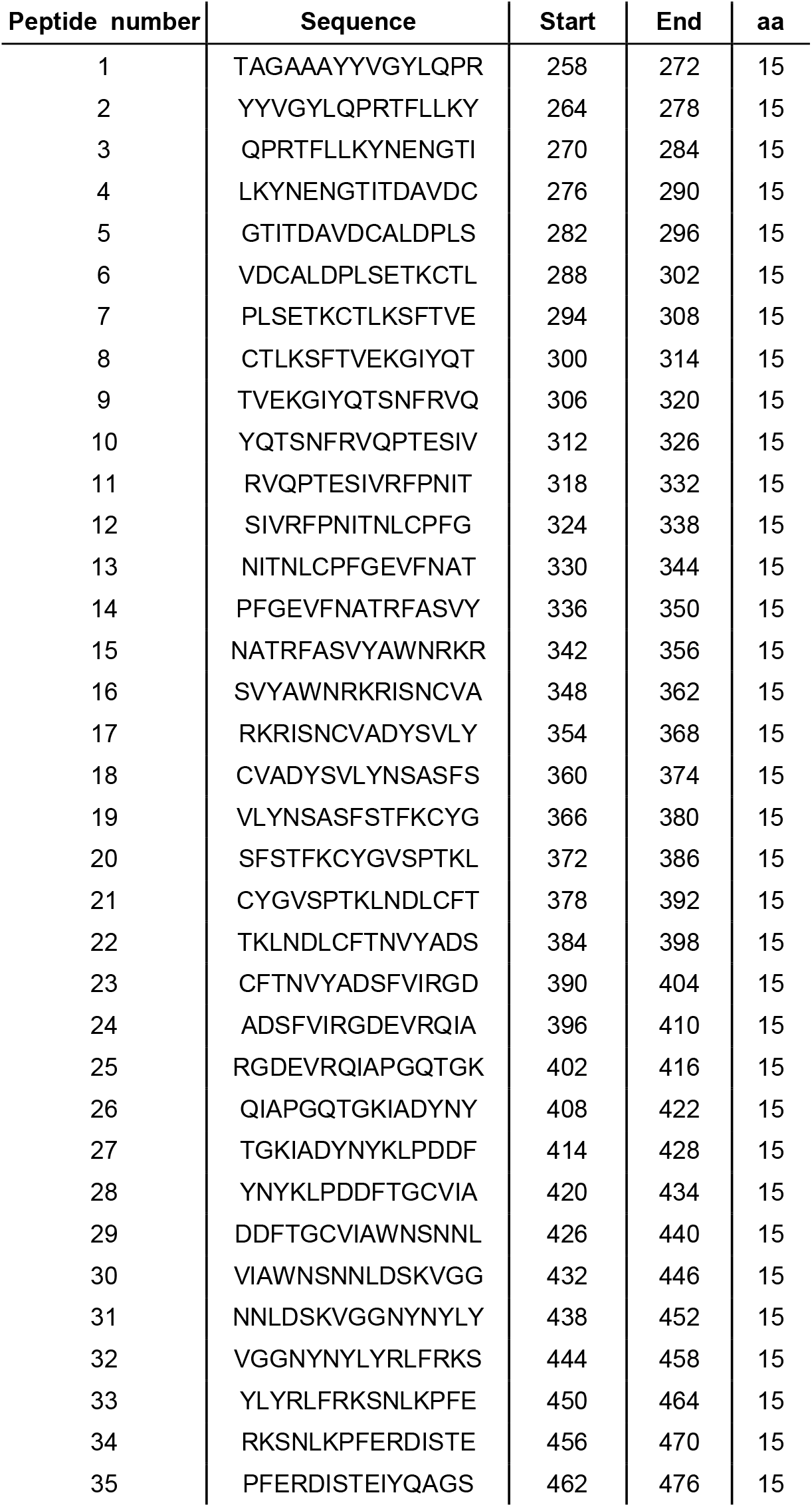
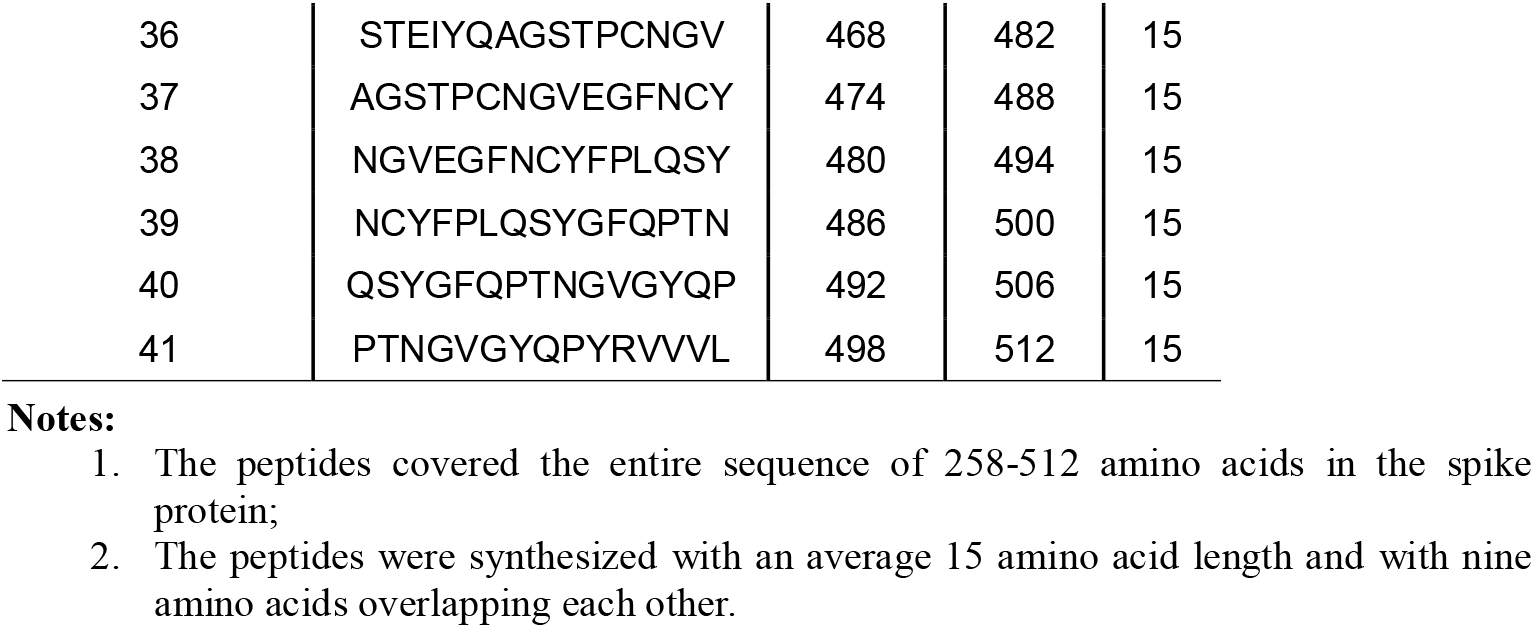
Overlapping Peptide Pool 1.

### Immunization

The mice were injected twice, with a two-week interval, via the intramuscular route (i.m.) with 25 μg pGX9501 expressing a synthetic, optimized sequence of the SARS-CoV-2 full-length spike glycoprotein^7^ and then electroporation was applied with the Cellectro2000 device. Serum samples and spleens were collected 14 days after the second immunization.

### IEDB analysis for SARS-CoV-2 MHC-I epitope identification

An explorative pannnel of SARS-CoV-2-derived epitopes with strongest predicted affinity to MHC Class I molecules was defined by Immune Epitope Database analysis (www.IEDB.org). The selection was based on internal predicitons using NetMHCpan Version EL4.1. all predicted epitopes with a percentile rank of <2 was selected for MHC-I processing analysis using MHC-NP methods. Stimultaneously, those eiptopes was analyzed in MHC-I immunogencity for checking whether the peptide sequence are consistent to the site preference of this allele. Combining results of the above three analysis methods, the peptide with a percentile rank of <0.5, a TAP total score of > -1 and a immunogencity score of >0 was picked up for the next ELIspot assay of T cell IFN-γ response.

### Cytotoxic lymphocyte (CTL) killing ability

A single-cell suspension of splenocytes from naïve syngeneic mice was diluted to 1.5*10^8^/ml in RPMI1640 containing 10% FBS and 2% penicillin and streptomycin and pulsed at 37°C with or without 5 ug/ml peptides as described previously^22^. After 4 h, eflour450 (eBioscience, 65-0842-85) at 5 mM (high concentration) was used to label peptide-pulsed cells at room temperature in the dark. The non-peptide-pulsed cells were labeled with a low concentration of eflour450 at 0.5 mM. After being rinsed three times with PBS, 4*10^6^ labeled and peptide-pulsed cells and an equal number of labeled non-peptide-pulsed cells were adoptively transferred by tail vein injections into mice that had previously been immunized. Six hours later, the percentage of labeled cells in spleens was detected with LSRFortessa flow cytometry (BD) and analyzed by FlowJo (TreeStar). The following formula calculated the specific cell lysis: Specific cell lysis ability = (1-(percentage of cells incubated with peptide/percentage of cells incubated without peptide)) *100%.

### IFN-γ ELIspot

Spleens were collected from mice individually into RPMI1640 media supplemented with 10% FBS (Gibco) and penicillin/streptomycin and processed into single-cell suspensions. ELIspot assays were performed using Mouse IFN-γ ELIspot plates (Dakewei Biotech Co., Ltd, 2210006). The ELIspot plates were repeat washed 5 times at RT with 100 μL of PBS per well and incubated with 200 μL of RPMI1640 10% FBS (R10) for 10 min before cell plating. Two hundred fifty thousand mouse splenocytes, CD4+, or CD8+ T cells were plated into each well and stimulated for 16 h with 15-mer peptides from the SARS CoV-2 Spike peptide pools that overlapped by nine amino acids as previously described^7^. Each peptide was present at a final concentration of 1 μg in 100 μl R10 per well. The spots were developed based on the manufacturer’s instructions. R10 and cell stimulation cocktails (Invitrogen) were used for negative and positive controls, respectively. Spots were scanned and quantified by AID ELIspot READER (AID, Germany). Spot-forming units (SFU) per million cells were calculated after subtracting the negative control wells.

### Statistical Analysis

The statistical analysis methods and sample sizes (n) are specified in the results section or figure legends for all quantitative data. All values are reported as means ± sem with the indicated sample size. No samples are excluded from the analysis. All relevant statistical tests are two-sided. P values less than 0.05 were considered statistically significant. All animal studies were performed with randomized animal selection. Statistics were performed using GraphPad Prism 7 software. In all data, *p<0.05, **p<0.01, ***p<0.001, and ****p<0.0001.

## RESULTS

### Strong CD8+ CTL epitope activity is embedded in an overlapping peptide pool 1 that covers the NTD and RBD region of the spike protein

When Balb/c mice were immunized twice with the pGX9501 DNA vaccine expressing the spike protein of SARS-CoV-2, a higher level of IFN-γ expression by splenocytes was often seen by the ELIspot assay when the cells were stimulated in vitro with spike peptide pool 1 (Table 1) compared with pool 2 (Figure 1A). In addition, when an in vivo CTL assay was done with identically immunized animals, the same peptide pool 1 gave a strong CTL response in vivo (Figure 1B), suggesting that MHC-I epitope(s) were present within pool 1.

**Figure 1.**
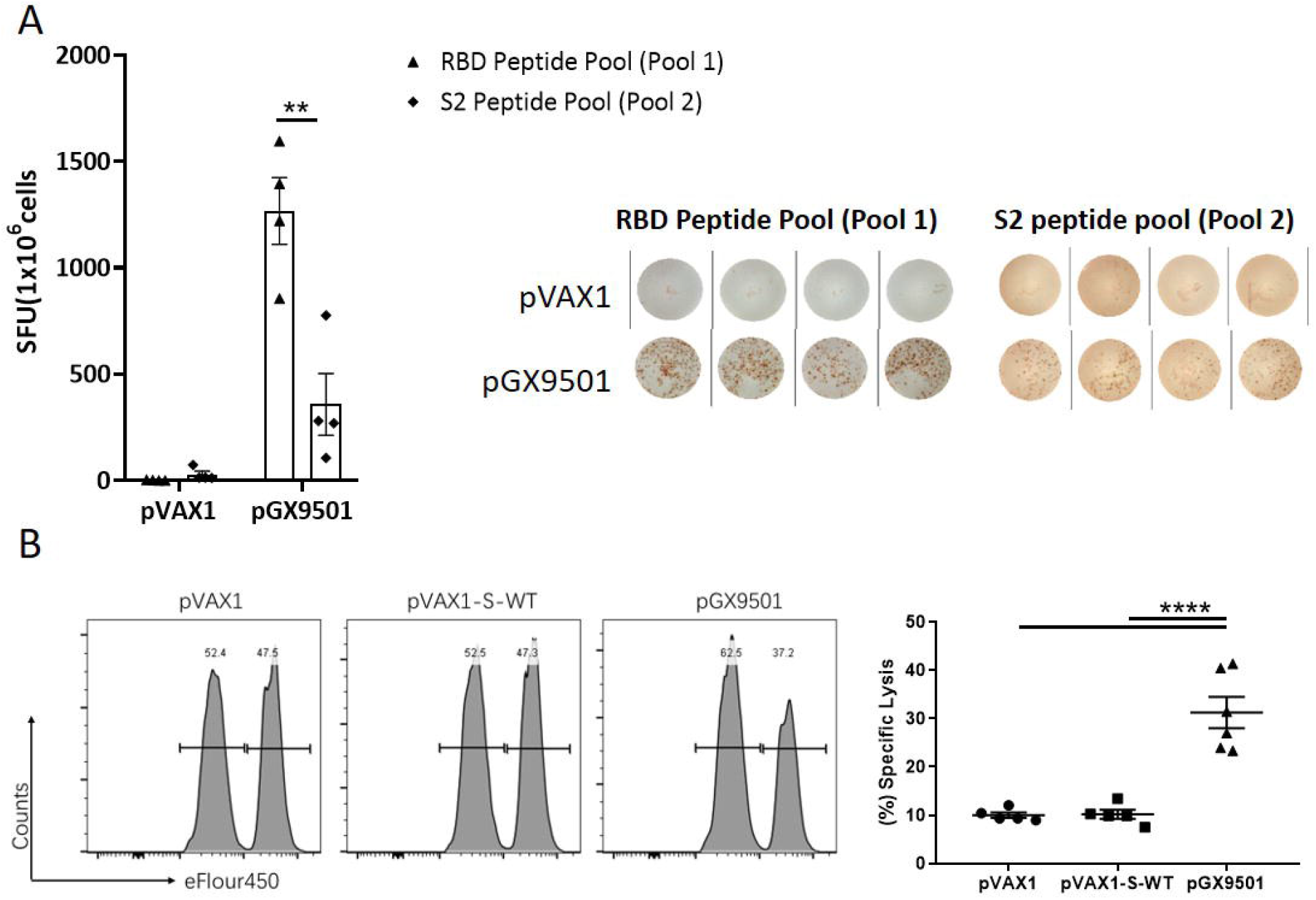
Peptide pool 1 Induced strong T cell responses in BALB/c mice post-administration of pGX9501. Balb/c mice (n = 5/group) were immunized twice two weeks apart with 25 μg pGX9501or pVAX1 (empty vector). T cell responses were analyzed on day 14 after the second injection. (A) Splenocytes were harvested and T cell responses were measured by IFN-γ ELIspot after stimulation for 20 h with overlapping peptide pools 1 or 2. (B) Antigen-specific cytotoxic lymphocyte (CTL) killing activity was evaluated by an in vivo CTL assay. Target cells at 4*10^6^/ml from naïve mice were peptide-pulsed with pool 1 then labeled with a high concentration of eFlour450 in vitro Control cells were non-peptide-pulsed cells and labeled with a low concentration of eFluor450.The cells were mixed and transferred i.v. into immunized mice. After 5 h, splenocytes were harvested and the intensity of eFlour450 peptide labeled target cells was compared with the non-peptide-labeled negative control cells by flow cytometry.

### Screening and identification of an MHC-I epitope in peptide pool 1

We placed the entire 41 peptide sequences from peptide pool 1 into the Immune Epitope Database analysis (IEDB) (http://www.iedb.org/) to seek critical epitopes. An evaluation method was established by integrating MHC-I binding prediction, MHC-I immunogenicity, and MHC natural processing (MHC-NP) prediction from three H-2d MHC-I alleles to improve the accuracy of prediction results (Table 2). In the H-2D^d^ allele, peptide 2 showed good MHC-I binding ability, immunogenicity, and TAP ability. In the H-2K^d^ allele, peptide 12 showed the strongest immunogenicity, and peptide 2 presented the most potent MHC-I binding ability and TAP ability among all peptides. In the H-2L^d^ allele, both peptide 2 and peptide 11 showed the strongest immunogenicity, while peptide 12 emerged as having the most potent TAP ability (Figure 2A&2B). Consequently, we used peptide 2, peptide 11, and peptide 12 to conduct an ELIspot assay of T cell IFN-γ response.

**Table 2.** MHC-I epitope analysis for Overlapping Peptide Pool 1.

| allele | peptide number | MHC-I binding | immunogenicity | Proteasome Score | TAP Score | MHC Score | Processing Score | TAP Total Score |
| --- | --- | --- | --- | --- | --- | --- | --- | --- |
| H-2-Dd | 21 | 0.01 | 0.03612 | 1.44 | 1.12 | -3.49 | 2.57 | -0.92 |
| H-2-Dd | 41 | 0.04 | 0.13706 | 1.74 | 0.4 | -2.56 | 2.14 | -0.42 |
| H-2-Dd | 2 | 0.05 | 0.1573 | 1.36 | 1.07 | -3.26 | 2.43 | -0.83 |
| H-2-Dd | 26 | 0.13 | -0.02676 | 1.38 | 1.27 | -4.51 | 2.65 | -1.86 |
| H-2-Dd | 9 | 0.19 | -0.11058 | 1.27 | 0.99 | -3.48 | 2.26 | -1.22 |
| H-2-Dd | 19 | 0.25 | 0.03263 | 1.33 | 1.04 | -3.45 | 2.37 | -1.08 |
| H-2-Dd | 12 | 0.29 | 0.1431 | 1.05 | 1.29 | -3.9 | 2.35 | -1.55 |
| H-2-Dd | 11 | 0.49 | 0.1386 | 1.36 | 1.21 | -4.02 | 2.57 | -1.45 |
| H-2-Dd | 32 | 0.62 | 0.0966 | 1.19 | 1.17 | -3.83 | 2.36 | -1.46 |
| H-2-Dd | 37 | 0.65 | 0.12191 | 1.38 | 1.09 | -3.76 | 2.47 | -1.29 |
| H-2-Dd | 14 | 0.67 | 0.08562 | 1.39 | 0.82 | -3.76 | 2.21 | -1.55 |
| H-2-Dd | 31 | 0.7 | 0.023 | 1.31 | 1.15 | -4.17 | 2.46 | -1.71 |
| H-2-Dd | 20 | 0.7 | -0.31841 | 1.47 | 1.3 | -4.6 | 2.77 | -1.83 |
| H-2-Dd | 27 | 0.89 | -0.11289 | 1.4 | 1.21 | -4.54 | 2.61 | -1.92 |
| H-2-Dd | 28 | 1.1 | -0.19576 | 0.98 | 1.12 | -4.22 | 2.1 | -2.12 |
| H-2-Dd | 24 | 1.2 | 0.12947 | 1.27 | 0.71 | -4.63 | 1.98 | -2.64 |
| H-2-Dd | 25 | 1.2 | -0.11559 | 1.43 | 0.15 | -4.39 | 1.58 | -2.81 |
| H-2-Dd | 18 | 1.2 | -0.22309 | 1.33 | 1.17 | -4.05 | 2.5 | -1.55 |
| H-2-Dd | 40 | 1.3 | 0.1256 | 1.38 | 1.34 | -4.5 | 2.71 | -1.79 |
| H-2-Dd | 1 | 1.3 | 0.06158 | 1.24 | 1.23 | -4.32 | 2.47 | -1.85 |
| H-2-Dd | 33 | 1.5 | -0.21085 | 1.03 | 1.24 | -4.23 | 2.27 | -1.96 |
| H-2-Dd | 29 | 1.6 | 0.05792 | 1.52 | 0.5 | -4.17 | 2.02 | -2.15 |
| H-2-Dd | 30 | 1.6 | 0.05792 | 1.35 | 0.46 | -4.17 | 1.8 | -2.36 |
| H-2-Dd | 39 | 1.6 | -0.15021 | 1.25 | 1.15 | -3.84 | 2.41 | -1.44 |
| H-2-Dd | 10 | 2 | 0.01977 | 1.1 | 0.24 | -4.23 | 1.34 | -2.89 |
| H-2-Kd | 2 | 0.01 | 0.06572 | 1.24 | 0.48 | -1.78 | 1.72 | -0.06 |
| H-2-Kd | 41 | 0.05 | 0.04196 | 1.74 | 0.46 | -3.43 | 2.2 | -1.22 |
| H-2-Kd | 9 | 0.06 | 0.12441 | 1 | 0.23 | -2.55 | 1.23 | -1.32 |
| H-2-Kd | 4 | 0.07 | 0.28634 | 1.34 | 0.37 | -2.34 | 1.71 | -0.63 |
| H-2-Kd | 3 | 0.07 | 0.05892 | 1.16 | 0.37 | -2.34 | 1.54 | -0.8 |
| H-2-Kd | 20 | 0.08 | 0.25644 | 1.45 | 0.49 | -2.65 | 1.93 | -0.71 |
| H-2-Kd | 21 | 0.08 | 0.05832 | 1.75 | 0.36 | -2.65 | 2.12 | -0.53 |
| H-2-Kd | 16 | 0.11 | 0.16858 | 1.36 | 0.4 | -2.32 | 1.77 | -0.55 |
| H-2-Kd | 40 | 0.19 | 0.0905 | 1.38 | 1.32 | -3.9 | 2.69 | -1.21 |
| H-2-Kd | 23 | 0.26 | 0.0573 | 1.31 | 0.44 | -3.18 | 1.75 | -1.43 |
| H-2-Kd | 24 | 0.61 | -0.0378 | 1.07 | 0.2 | -3.02 | 1.27 | -1.75 |
| H-2-Kd | 19 | 0.68 | 0.0279 | 1.33 | 1.2 | -3.27 | 2.52 | -0.75 |
| H-2-Kd | 10 | 0.79 | 0.034 | 1.1 | 0.23 | -3.35 | 1.33 | -2.02 |
| H-2-Kd | 12 | 0.92 | 0.34063 | 1.45 | 0.59 | -3.13 | 2.03 | -1.1 |
| H-2-Kd | 1 | 0.96 | -0.04018 | 1.24 | 1.29 | -4.51 | 2.53 | -1.98 |
| H-2-Kd | 32 | 0.99 | 0.13255 | 1.19 | 1.18 | -3.56 | 2.37 | -1.19 |
| H-2-Kd | 39 | 1.1 | 0.0801 | 1.25 | 1.31 | -3.16 | 2.56 | -0.6 |
| H-2-Kd | 31 | 1.2 | 0.1811 | 1.48 | 0.48 | -3.32 | 1.96 | -1.36 |
| H-2-Kd | 29 | 2 | 0.07062 | 1.52 | 0.5 | -3.5 | 2.02 | -1.48 |
| H-2-Kd | 30 | 2 | 0.07062 | 1.35 | 0.46 | -3.5 | 1.8 | -1.7 |
| H-2-Kd | 35 | 2 | -0.15381 | 1.42 | 1.16 | -4.52 | 2.57 | -1.95 |
| H-2-Ld | 11 | 0.06 | 0.30371 | 1.36 | 0.98 | -3.5 | 2.34 | -1.16 |
| H-2-Ld | 41 | 0.12 | -0.07228 | 1.74 | 0.35 | -3.54 | 2.09 | -1.45 |
| H-2-Ld | 21 | 0.2 | 0.05832 | 1.44 | 0.99 | -3.65 | 2.44 | -1.21 |
| H-2-Ld | 39 | 0.27 | -0.19696 | 1.25 | 0.91 | -3 | 2.16 | -0.84 |
| H-2-Ld | 12 | 0.28 | 0.09851 | 1.05 | 0.94 | -2.75 | 2 | -0.75 |
| H-2-Ld | 37 | 0.44 | 0.0801 | 1.38 | 0.99 | -4.01 | 2.37 | -1.65 |
| H-2-Ld | 2 | 0.54 | 0.28634 | 1.41 | 1.19 | -3.97 | 2.6 | -1.38 |
| H-2-Ld | 3 | 0.57 | -0.08994 | 1.52 | 1.13 | -3.97 | 2.65 | -1.32 |
| H-2-Ld | 27 | 0.77 | 0.0573 | 1.4 | 1.21 | -4.39 | 2.61 | -1.78 |
| H-2-Ld | 28 | 0.77 | 0.03448 | 0.98 | 1.12 | -4.18 | 2.1 | -2.07 |
| H-2-Ld | 19 | 0.95 | 0.11915 | 1.53 | 1.42 | -4.32 | 2.95 | -1.37 |
| H-2-Ld | 18 | 1.1 | -0.22669 | 1.33 | 1.17 | -3.82 | 2.5 | -1.32 |
| H-2-Ld | 34 | 1.9 | 0.1811 | 0.91 | 1.16 | -4.58 | 2.07 | -2.5 |
| H-2-Ld | 9 | 1.9 | -0.08994 | 1.27 | 0.99 | -4.13 | 2.26 | -1.87 |
| H-2-Ld | 38 | 2 | -0.19696 | 0.99 | 0.28 | -2.8 | 1.27 | -1.53 |

Notes:
1. MHC-I binding score was between 0 and 2. <0.5 strong binder, 0.5-2 weak binder, >2 non-binder. 2. A high Immunogenicity score indicates that the degree of the peptide conformity to sequence preference was good. 3. The higher the TAP total score, the higher the likelihood that the peptide will be presented after being swallowed by DCs.

**Figure 2.**
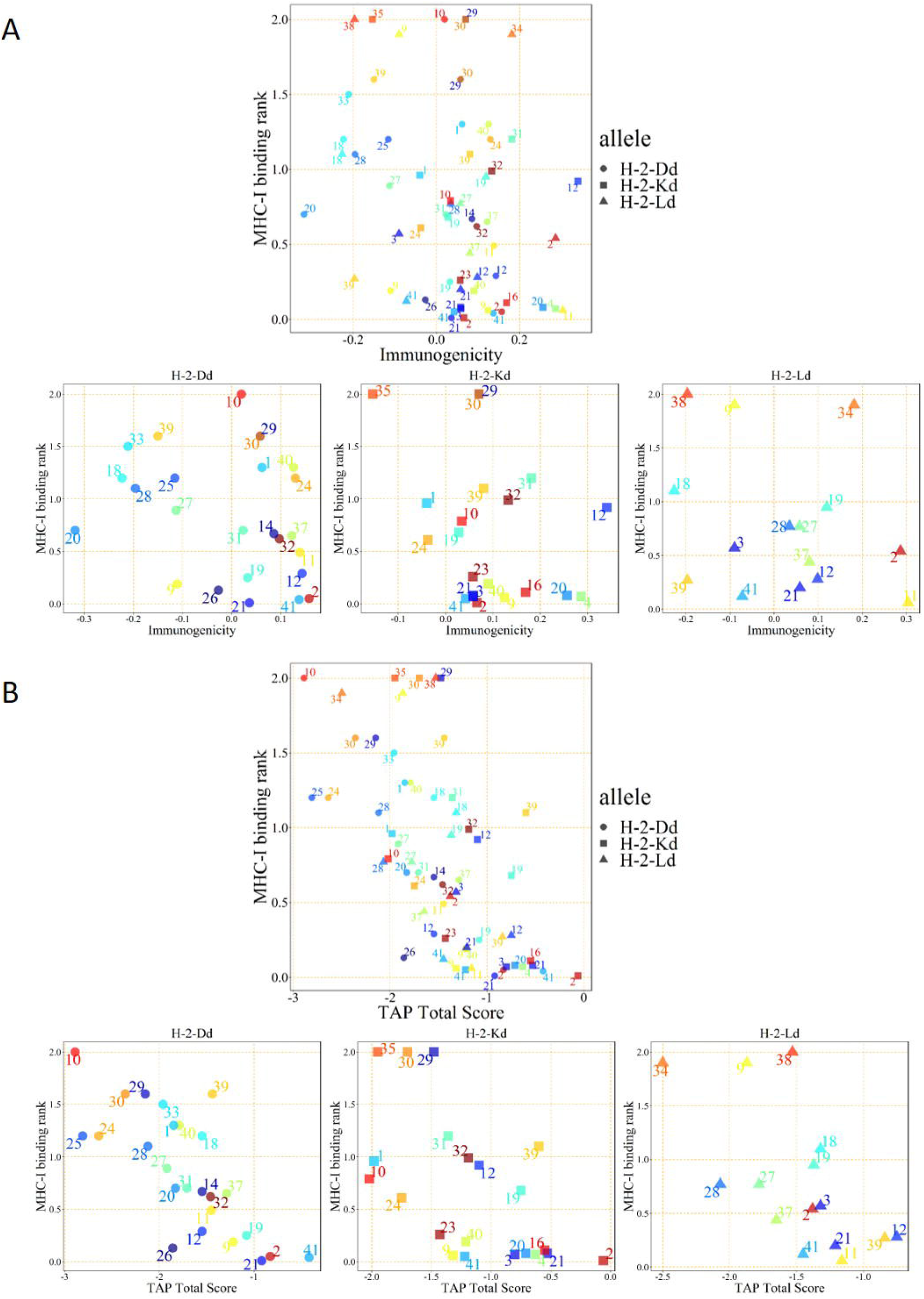
IEDB prediction scores of peptides in pool 1. CTL epitope peptides were screened by integrating MHC-I binding prediction, MHC-I immunogenicity (A), and MHC-NP (B) prediction from three H-2d MHC-I alleles.

As shown in Figure 3A, peptide 2 from pool 1 presented the best stimulation to induce the IFN-γ secretion compared to the other two selected peptides. Thus, peptide 2, consisting of 15 amino acids, stimulated CD8+ T cells via MHC-I or/and CD4+ T cells via MHC-II. To identify which T cell type was stimulated by peptide 2, purified CD4+ T cells or CD8+ T cells were used (sFigure 1). Peptide 2 stimulated CD8+ T cells but not CD4+ cells, indicating that it can only be presented by MHC-I (Figure 3B). To further investigate its sequence specificity, we mutated several predicted anchor amino acids of peptide 2 according to the preferences of the H-2d MHC-I allele^11-14^. The mutated peptide 2 had a low MHC-I binding score in the IEDB prediction (Table 3) and showed a significantly reduced ability to stimulate IFN-γ secretion by CD8+ T cells (Figure 3C).

**Table 3.**
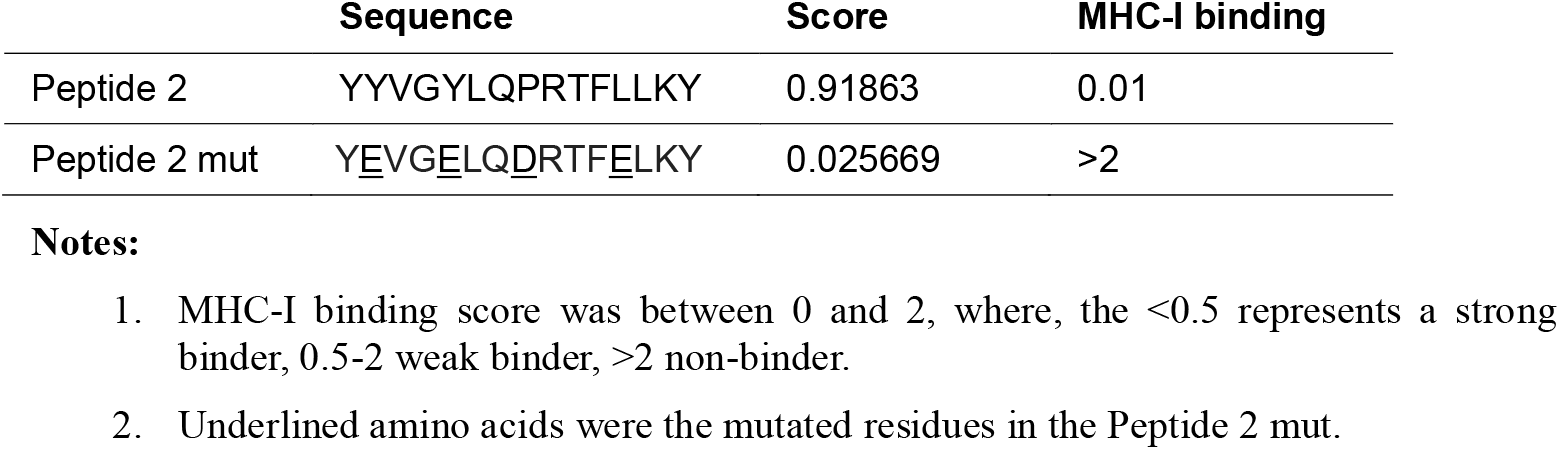
Prediction scores of peptide 2 and mutated peptide 2.

**Figure 3.**
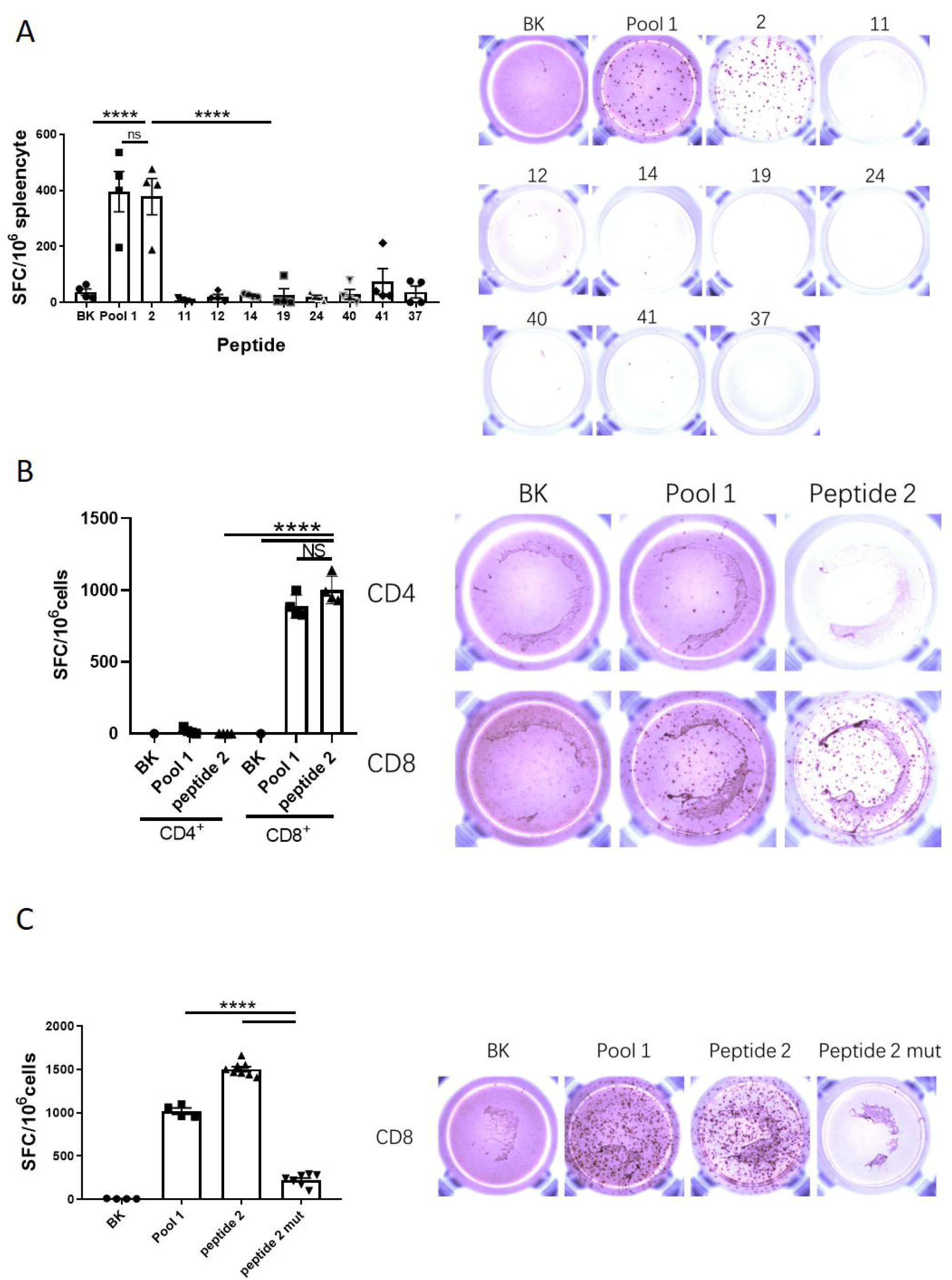
Peptide 2 is identified as the CD8+ CTL epitope. BALB/c mice were immunized with the pGX9501. Splenocytes were obtained and specific T-cell induction was analyzed with IFN-γ ELIspot, using stimulation with the indicated peptides. (A) Splenocytes, (B) CD4+, and (C) CD8+ T cells..

### Analysis of peptide 2 epitope conservation and HLA distribution

We used the peptide 2 sequence, YYVGYLQPRTFLLKY (amino acid 264-278), to compare with current posted SARS-CoV-2 variants including variant-of-interest (VOI) and variant-of-concern (VOC) published by WHO including the lastest ο variant and observed that this sequence is highly conserved (Figure 4A), and located at the end of the NTD of Spike protein and upstream of the RBD (Figure 4B). The highly conserved epitopic sequence provides a useful tool for evaluating the CD8+ T cell-mediated responses to vaccination in animals and humans.

**Figure 4.**
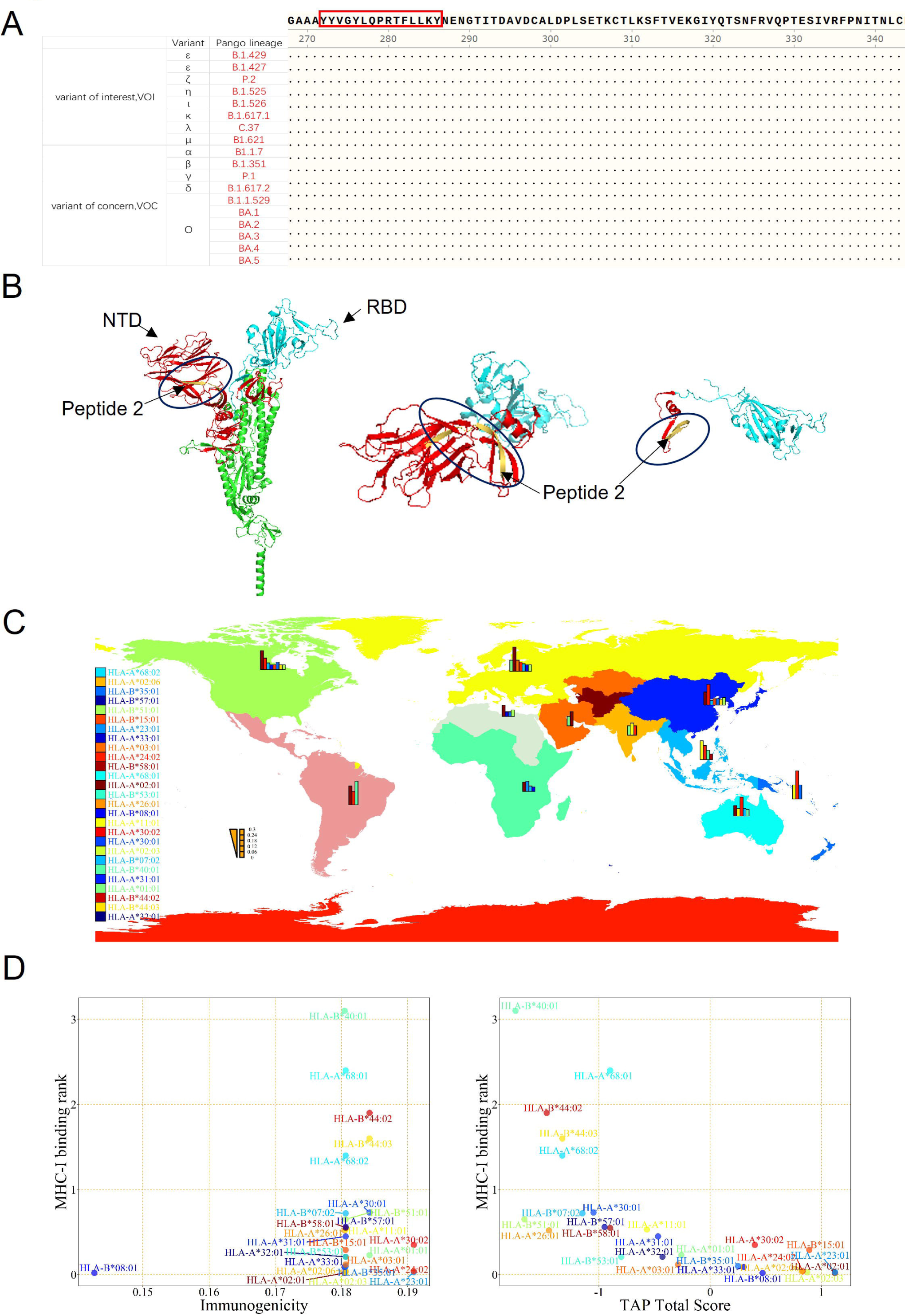
Conserved Sequence & MHC-I HLA Analysis of peptide 2. (A) The sequence of Peptide 2 was highly conserved in 16 SARS-CoV-2 variants (including 5 subvariants of omicron) that have been identified as the variant of interest and the variant of concern published by WHO. (B) Position of Peptide 2 (yellow marked segment) in the stereoscopic structure of the spike protein. (C) The global distribution of HLA alleles. (D) Analysis of Peptide 2 by integration of MHC-I binding prediction, MHC-I immunogenicity, and MHC-NP prediction from the HLA alleles.

Since MHC-I-biased expression patterns in different populations globally are diversified and variable, the peptide 2 sequence that can be recognized by one population may not be recognized by others. To investigate if peptide 2 sequence could be recognized by most populations worldwide, we performed an HLA allele analysis for different regions to assess binding to one or more of the 27 prevalent MHC-I molecules, including HLA-A (01:01, 02:01, 02:03, 02:06, 03:01, 11:01, 23:01, 24:02, 26:01, 30:01, 30:02, 31:01, 32:01, 33:01, 68:01, 68:02), and HLA-B (07:02, 08:01, 15:01, 35:01, 40:01, 44:02, 44:03, 51:01, 53:01, 57:01, 58:01), as shown in Figure 4C & Table 4. The frequency by HLAs was calculated by the online analysis tool at http://www.allelefrequencies.net/. We also evaluated the MHC-I binding ability, immunogenicity, and TAP potential of peptide 2 on different HLA alleles by IEDB (Figure 4D & Table 5). The results indicated that peptide 2 could be recognizable by the HLA-A*02:01 allele (most in Europe and America), HLA-B*08:01 allele (in Europe and Australia), HLA-A*23:01 allele (in North Africa and Sub-Saharan Africa), HLA-A*02:03 allele (in Southeast Asia), HLA-A*24:02 allele (in Oceania), HLA-A*02:06 allele (in North America, North-East Asia, and Oceania), HLA-A*33:01 allele (in China and Pakistan), HLA-B*35:01 allele (in Oceania), and HLA-A*03:01 allele (in Europe). These findings suggest that peptide 2 could be well recognized by the most frequent HLA alleles of the worldwide population and can therefore be considered to be promiscuous.

**Table 4.** Geographic Distribution of HLA allele.

| **Continent** | **Allele** | **Frequency** | **Allele** | **Frequency** |
| --- | --- | --- | --- | --- |
| Australia | HLA-A*24:02 | 0.2 | HLA-B*07:02 | 0.08 |
|  | HLA-A*02:01 | 0.11 | HLA-B*40:01 | 0.07 |
|  | HLA-A*11:01 | 0.08 |  |  |
| Europe | HLA-A*02:01 | 0.26 | HLA-B*07:02 | 0.08 |
|  | HLA-A*01:01 | 0.12 | HLA-B*08:01 | 0.07 |
|  | HLA-A*03:01 | 0.12 | HLA-B*51:01 | 0.07 |
|  | HLA-A*24:02 | 0.1 |  |  |
| North Africa | HLA-A*02:01 | 0.12 | HLA-B*51:01 | 0.07 |
|  |  |  | HLA-B*08:01 | 0.05 |
|  |  |  | HLA-B*35:01 | 0.05 |
| North America | HLA-A*02:01 | 0.2 | HLA-B*35:01 | 0.08 |
|  | HLA-A*24:02 | 0.12 | HLA-B*07:02 | 0.07 |
|  |  |  | HLA-B*08:01 | 0.05 |
|  |  |  | HLA-B*15:01 | 0.05 |
|  |  |  | HLA-B*44:03 | 0.05 |
|  |  |  | HLA-B*51:01 | 0.05 |
| North-East Asia | HLA-A*24:02 | 0.22 | HLA-B*51:01 | 0.08 |
|  | HLA-A*02:01 | 0.14 | HLA-B*35:01 | 0.07 |
|  |  |  | HLA-B*15:01 | 0.07 |
|  |  |  | HLA-B*44:03 | 0.06 |
|  |  |  | HLA-B*07:02 | 0.05 |
| Oceania | HLA-A*24:02 | 0.3 | HLA-B*35:01 | 0.15 |
|  | HLA-A*11:01 | 0.15 |  |  |
| South and Central America | HLA-A*02:01 | 0.2 | HLA-B*40:01 | 0.25 |
|  | HLA-A*24:02 | 0.14 |  |  |
| South Asia | HLA-A*11:01 | 0.13 |  |  |
|  | HLA-A*01:01 | 0.1 |  |  |
|  | HLA-A*24:02 | 0.1 |  |  |
| South-East Asia | HLA-A*11:01 | 0.2 | HLA-B*40:01 | 0.1 |
|  | HLA-A*24:02 | 0.15 | HLA-B*58:01 | 0.06 |
| Sub-Saharan Africa | HLA-A*23:01 | 0.11 | HLA-B*07:02 | 0.06 |
|  | HLA-A*02:01 | 0.1 | HLA-B*08:01 | 0.05 |
| Western Asia | HLA-A*02:01 | 0.15 |  |  |
|  | HLA-A*01:01 | 0.1 |  |  |

**Table 5.**
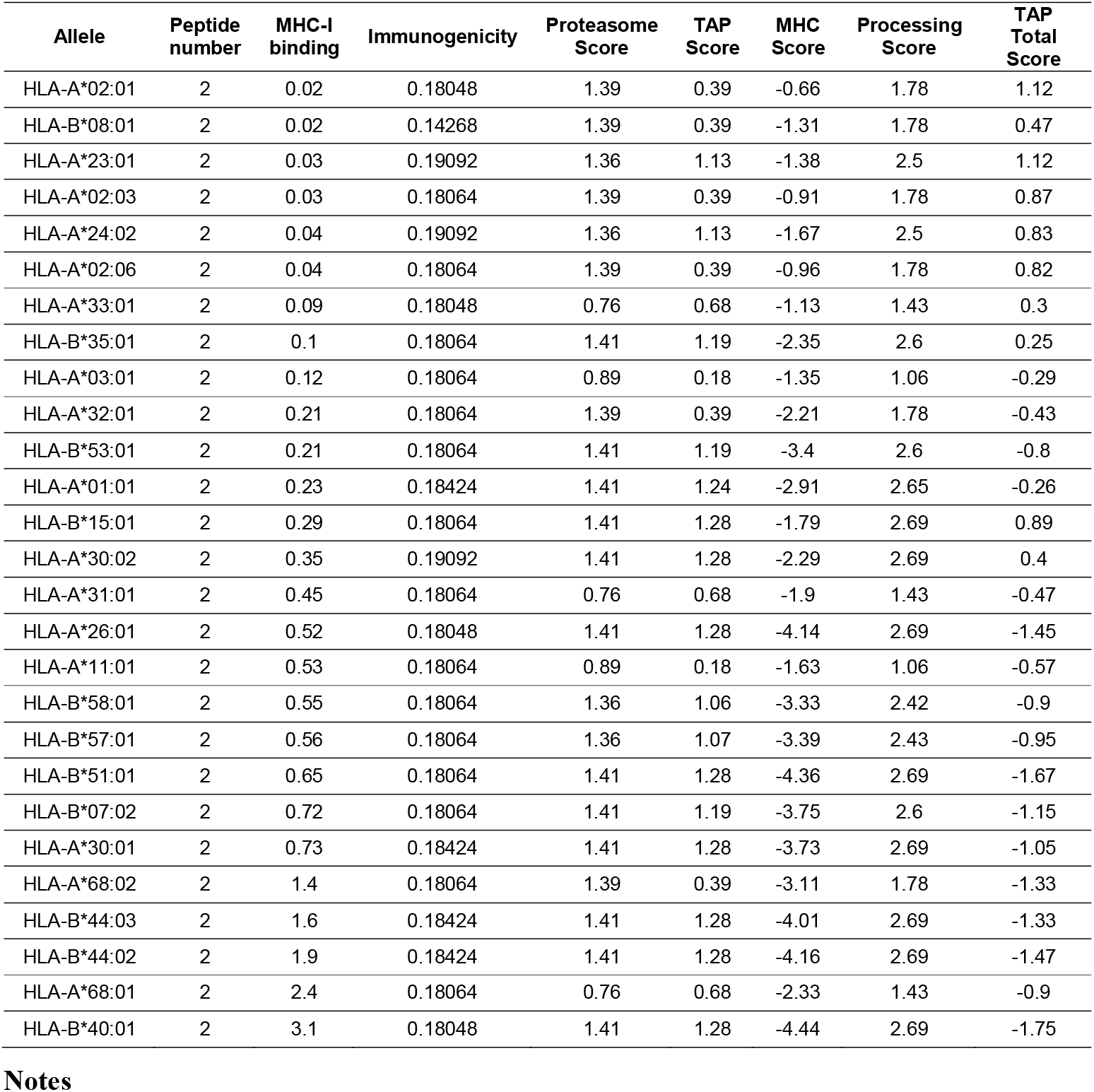
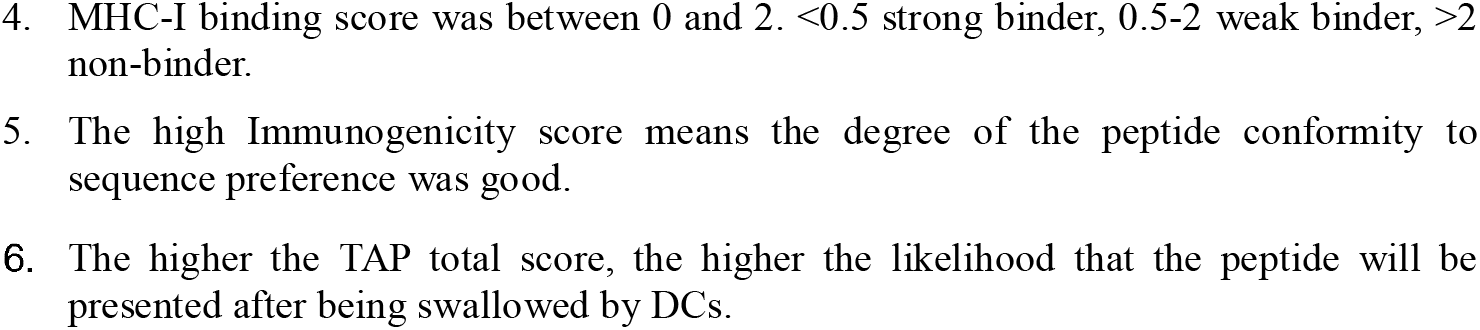
Human MHC-I epitope analysis for peptide 2.

## DISCUSSION

In this study, we have defined and characterized a potential CTL epitope conserved and established a BALB/c mice model to evaluate CTL responses to the spike protein of SARS-CoV-2. Furthermore, we found the peptide 2 may be well recognized by HLA alleles in most populations worldwide.

In recent studies, CD8+ T cell immunity was found to make major contributions to the protective efficacy of SARS-CoV-2 vaccines^15-17^. Additionally, lymphopenia was more accentuated in symptomatic COVID-19 patients with pneumonia than in those without pneumonia, consistent with T cell immunity playing a protective role in pre-existing immunity against SARS-CoV-2^17-19^. However, the role of T cell immunity in the pathology of COVID-19 has not been clarified^20^ and awaits further definition of T cell epitopes and their functions. Engineered improvements in the peptide epitopes recognized by CTL have been an essential feature of the advancements made in evaluating the cellular immunity induced by vaccines. Our work has provided a tool to monitor virus-specific CD8+ T cells and assess the contribution of CTLs to the control and the elimination of the virus.

The SARS-CoV-2 virus was found to mutate rapidly, the development of vaccines that could protect people from different variants was seen to be urgent. The neutralizing antibodies induced by vaccines were found to have variable efficacies against the different SARS-CoV-2 variants and efficacies declined over time, whereas the protection represented by CD8^+^ T cell immunity remained unchanged^15^. We showed here that one particular peptide from the spike protein, peptide 2 is highly conserved among variants and could be well presented by MHC-1 of all of the HLA alleles that are most frequent worldwide. Thus, the conserved peptide 2 was found suitable for developing peptide vaccines to induce cellular immunity against different variants of SARS-CoV-2, including variant-of-interest (VOI) and variant-of-concern (VOC). A recent study confirmed that peptide 2 probably has a strong cell-mediated immunological function in man; a 9-mer (YLQPRTFLL) peptide that was overlapped by peptide 2 was able to induce a high level of IFN-γ expression from PBMCs of patients who had recovered from COVID-19 and carried the HLA-A*02:01 allele^21^.

In conclusion, our study utilized web-based tools to predict human MHC-I epitopes although peptide 2 (YYVGYLQPRTFLLKY) did not give the strongest total TAP score in the prediction, the simulations indicated particularly robust antigen-specific IFN-γ-expressing CD8+ T responses overall compared to the other predicted epitope sequences. This epitope sequence is highly conservative among currently known SARS-CoV-2 variants and recognizable by most world populations. This critical MHC-I epitope sequence is located at the end of NTD of the spike protein; it can be used to assess CMI induced by a COVID-19 vaccination; it may be strategically incorporated into vaccine designs to enhance the prospect of viral elimination by vaccination.

## ACKNOWLEDGMENT

This work was supported by the Chinese National Natural Science Foundation (81991492 and 82041039) and National Key R&D Program of Chinese Ministry of Science & Technology (2018YFC0840402) to B.Wang. We wish to thank Ms Yiwei Zhong at Shanghai Medical College of Fudan University for her techical supports..

## AUTHOR CONTRIBUTIONS

BW and SJ conceived, designed the study and drafted manuscript. BW supervised the study. SJ, SW, XG, JH, YD and ZZ performed the experiments. SJ, YH, GZ and AC analyzed the data. All authors reviewed the manuscript.

