## supplementary figure 1 for "Identification of a promiscuous conserved CTL epitope within the SARS-CoV-2 spike protein"

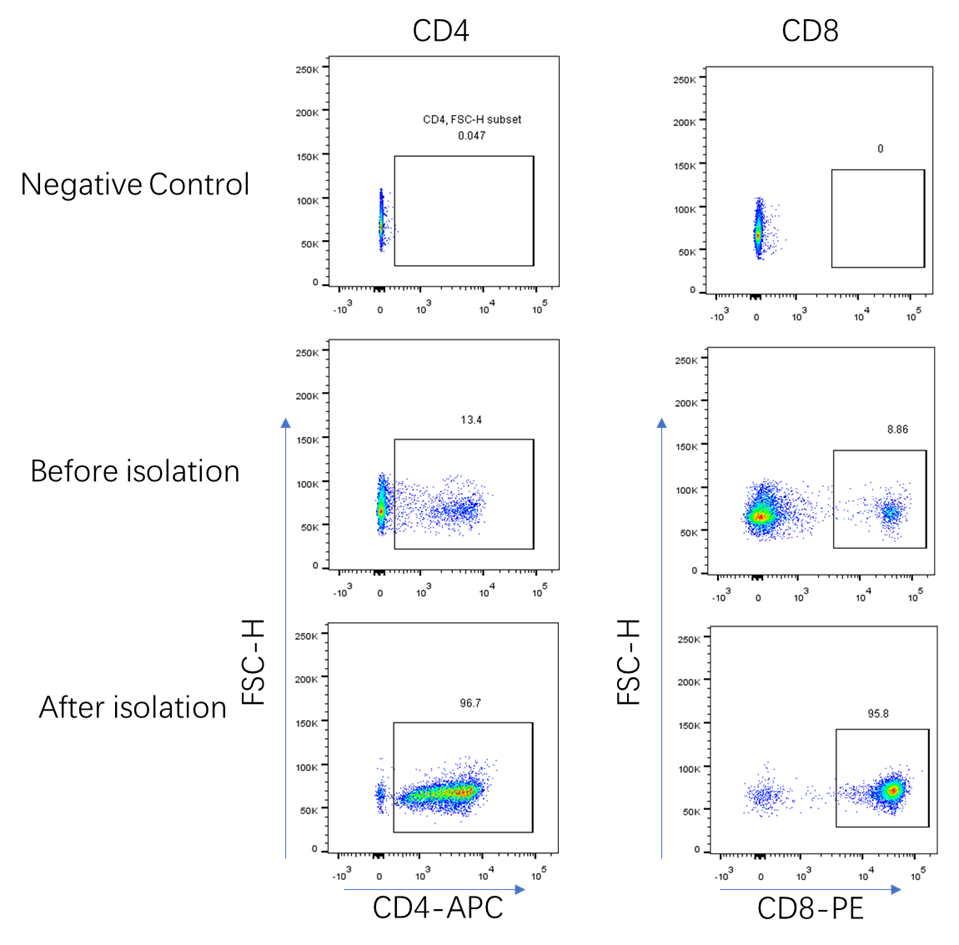

**sFigure1. Cell separation and analysis by flow cytometry.**

Separation of CD4+ T cells and CD8+ T cells from Balb/c mouse spleen.

Splenocytes were collected from mice immunized with pGX9501 and CD4-positive and CD8-positive T cells were separated by magnetic tag. The splenocytes and T cells were stained with CD4-APC and CD8-PE and detected with LSRFortessa flow cytometry (BD) and analyzed by FlowJo (TreeStar). Gates were set on lymphocytes (FSC/SSC).
